# Transcriptome and regulatory maps of decidua-derived stromal cells inform gene discovery in preterm birth

**DOI:** 10.1101/2020.04.06.017079

**Authors:** Noboru Sakabe, Ivy Aneas, Nicholas Knoblauch, Debora R. Sobreira, Nicole Clark, Cristina Paz, Cynthia Horth, Ryan Ziffra, Harjot Kaur, Xiao Liu, Rebecca Anderson, Jean Morrison, Virginia C. Cheung, Chad Grotegut, Timothy E. Reddy, Bo Jacobsson, Mikko Hallman, Kari Teramo, Amy Murtha, John Kessler, William Grobman, Ge Zhang, Louis J. Muglia, Sarosh Rana, Vincent J. Lynch, Gregory E. Crawford, Carole Ober, Xin He, Marcelo A. Nóbrega

## Abstract

While a genetic component of preterm birth (PTB) has long been recognized and recently mapped by genome-wide association studies (GWAS), the molecular determinants underlying PTB remain elusive. This stems in part from an incomplete availability of comprehensive functional genomic annotations in human cell types relevant to pregnancy and PTB. Here, we generated extensive transcriptional and chromatin annotations of cultured primary decidua-derived mesenchymal stromal/stem cells (MSCs) and *in vitro* differentiated decidual stromal cells (DSCs) and developed a computational framework to integrate these functional annotations with results from a GWAS of gestational duration in 56,384 women. This resulted in a significant enrichment of heritability estimates in functional noncoding regions in stromal cells, as well as in the discovery of additional loci associated with gestational duration and target genes of associated loci. Our strategy illustrates how systematic functional annotations in pregnancy-relevant cell types aid in the experimental follow-up of GWAS for PTB and, likely, other pregnancy-related conditions.

## Introduction

Spontaneous preterm birth (PTB), defined as spontaneous labor and birth before 37 weeks of gestation, is associated with significant infant mortality and morbidity, as well as long-term health consequences into adulthood^1^. A genetic component to PTB has long been recognized but the significant role of environmental factors and the etiologic heterogeneity of birth before 37 weeks^2-4^ has made it challenging to discover genetic associations and causal genes. For example, recent genome-wide association studies (GWASs) of gestational duration in 43,568 women (3,331 with a preterm delivery)^5^ and in 84,689 infants (4,775 born preterm)^6^ reported six and one genome-wide significant associations, respectively, with gestational duration considered as a continuous variable. Three loci were also associated with PTB (defined as a categorical variable of birth) in the maternal GWAS^5^, but no loci were associated with PTB in the infant GWAS^6^. These studies highlight the challenges of such complex and multifactorial phenotypes, and the need for additional approaches to facilitate discovery of genes contributing to gestational duration and PTB.

Integrating GWAS results with genomic and epigenomic annotations is a promising approach for assigning function to variants discovered by GWAS, as well as for identifying additional associations that do not reach stringent genome-wide significance threshold^7,8^. While large consortia (e.g., ENCODE^9^, GTEx^10^, Roadmap Epigenomics^11^) have generated annotations of putative functional elements and genetic variants for many human cell types and tissues, there is a remarkable absence in these databases for the cell types and tissues that are relevant to pregnancy in general and to PTB in particular. Because the regulation of transcription has strong cell type-specific components, and because annotations in disease-relevant tissues or cells tend to be most enriched among GWAS signals for those specific diseases^10,12^, follow-up studies of GWASs of pregnancy-associated conditions have been disadvantaged compared to most other complex diseases due to the paucity of functional annotations in cells relevant to pregnancy. To fill this gap in knowledge, we characterized the transcriptional and chromatin landscapes of cultured mesenchymal stromal/stem cells (MSCs) collected from human placental membranes and decidualized MSCs, also known as decidual stromal cells or DSCs. These cells play critical roles in promoting successful pregnancy, interfacing with fetal cells throughout pregnancy, and the timing of birth^13^. We then built a computational framework that integrated these decidua-derived stromal cell annotations with the results of a large GWAS of gestational duration to facilitate discovery of PTB genes.

This integrated analysis revealed a significant enrichment of heritability estimates for gestational duration in decidua-derived stromal cell genomic regions marked by open chromatin or histone marks. Leveraging those functional annotations in a Bayesian statistical framework, we discovered additional loci associated with gestational duration and improved fine mapping in regions associated with gestational duration. Finally, using promoter capture Hi-C (pcHi-C), we linked functionally-annotated gestational age-associated variants to their putative target genes. More generally, these functional annotations should prove a valuable resource for studying other pregnancy-related conditions, such as preeclampsia and recurrent miscarriage, as well as conditions associated with endometrial dysfunction, such as endometriosis and infertility.

## Results

### Generation of transcriptome and epigenome maps of untreated (MSC) and in vitro differentiated decidual stromal cells (DSC)

Decidualization is the process of transformation of endometrial MSCs into DSCs that is induced by progesterone production that begins during the luteal phase of the menstrual cycle and then increases throughout pregnancy when successful implantation occurs (reviewed in ^14^). Using progesterone and estrogen or cyclic AMP to induce decidualization of MSCs in culture has been used in cells derived from endometrial biopsies in non-pregnant women to characterize their transcriptomes and epigenomes and to identify genes and molecular pathways involved in this process^15-20^.

Because obtaining endometrial cells in non-pregnant women through biopsies requires an invasive procedure that carries some risk and MSCs can also be obtained from human placentas^21-23^, we isolated these cells from the decidua parietalis of three women who had delivered at term and established one primary MSC line from each to model the process of decidualization (see Materials and Methods). Briefly, cells were treated with medroxyprogesterone acetate (MPA) and cAMP for 48 hours and a paired set of untreated samples was cultured in parallel for 48 hours. Three replicates of treated/untreated sets of each cell line were studied to assess experimental variability in the 2 conditions. Each of the 18 samples (3 individual lines x 3 replicates x 2 conditions) were assayed to generate transcriptome (RNA-seq), open chromatin (ATAC-seq) and histone modification (ChIP-seq) maps. A summary of those data is shown in Table 1 and a representative example of the full set of annotations for one primary cell line is shown in Figure 1. The number of reads generated for each sample in each condition and other descriptive data are provided in Supplementary File S1.

**Table 1.** Summary of ChIP-seq, ATAC-seq, and RNA-seq data: number of peaks, expressed genes and differentially expressed genes

|  | Untreated (MSC) |  |  | Decidualized (DSC) |  |  | Differential |  |
| --- | --- | --- | --- | --- | --- | --- | --- | --- |
|  | Cell line 1 | Cell line 2 | Cell line 3 | Cell line 1 | Cell line 2 | Cell line 3 | Up in decidualized | Down in decidualized |
| H3K27ac | 116,725 | 109,608 | 128,770 | 115,548 | 117,617 | 117,833 | 15,370 | 11,544 |
| H3K4me1 | 204,756 | 187,740 | 189,959 | 190,495 | 189,419 | 190,151 | 13,970 | 5,203 |
| H3K4me3 | 69,708 | 50,298 | 63,838 | 81,663 | 45,707 | 65,919 | 1,883 | 368 |
| ATAC-seq | 164,755 | 177,877 | 149,396 | 158,142 | 107,200 | 110,927 | 5,562 | 2,395 |
| RNA-seq | 13,168 | 13,145 | 13,079 | 12,913 | 13,096 | 12,923 | 502 | 633 |

**Figure 1.**
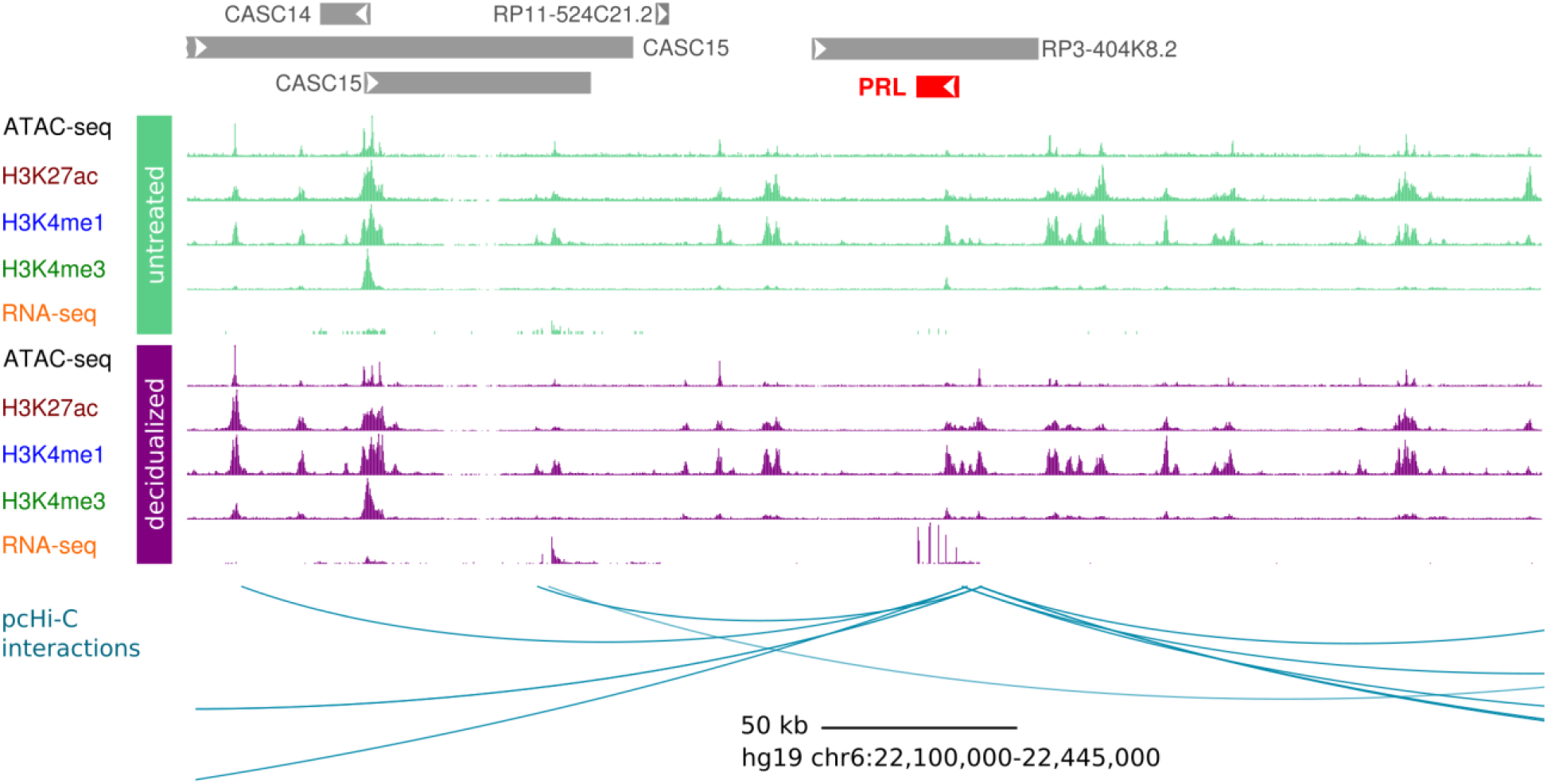
Schematic of RNA-seq, ATAC-seq, ChIP-seq, and pcHi-C maps centered on the prolactin (*PRL*) gene, as an example. Each histone modification and RNA-seq track shows read counts per base pair for each experiment. The pcHi-C signal track shows the number of reads per *Mbo*I restriction fragment. Arcs in the pcHi-C interactions track show significant interactions between the promoter of the *PRL* (prolactin) gene and putative distal regulatory elements identified with pcHi-C. Pooled data (3 replicates) for one cell line are shown for untreated cells (MSCs, in green) and decidualized cells (DSCs, in purple). pcHi-C data were generated in a fourth cell line that was decidualized.

### Robust gene expression changes occur in decidualized stromal cells

Analysis of the RNA-seq data using DESeq2^24^ revealed 1,135 differentially expressed genes after decidualization (Table 1). Genes with decreased expression after 48 hours of treatment were highly enriched for cell cycle genes (Supplementary File S2), consistent with observations from endometrial biopsies from non-pregnant women that decidualization is associated with cell cycle arrest^18,25^. Genes with increased expression after treatment were enriched for insulin-related terms, also consistent with previous results from endometrial biopsies^25^, and for glucose metabolism^17^.

### Identification of regulatory elements associated with decidualization

To identify putative regulatory elements in MSCs and DSCs, we assayed H3K27ac, H3K4me1 and H3K4me3 histone modifications, which are markers of active enhancers, poised enhancers, and active promoters, respectively (reviewed in ^26^). We also employed ATAC-seq to identify open chromatin regions to complement ChIP-seq data. To identify regulatory regions that might be altered in response to, and potentially regulate decidualization, we compared read counts of ATAC-seq and ChIP-seq peaks in untreated and decidualized cells, revealing tens of thousands of regions that differed between untreated and treated samples (Table 1). The majority of differential peaks were marked with H3K27Ac and H3K4me1, indicating that the epigenetic changes underlying alterations in gene expression during decidualization predominantly occur in distant regulatory elements, such as enhancers.

We observed a moderate degree of overlap between the differential peaks across ATAC-seq and ChIP-seq data, with the two enhancer marks, H3K27ac and H3K4me1, showing the most overlap (Figure 2a). Additionally, putative regulatory regions that showed chromatin changes in response to decidualization were associated with genes whose expression also changed in response to decidualization (Figure 2b). Regulatory regions with increased read counts clustered around genes that were more highly expressed after decidualization, indicating increased chromatin accessibility or activation of enhancers of those genes. Conversely, genes that were more lowly expressed after decidualization were enriched for enhancers that became less accessible or active. These observations indicate that the differential peaks of open chromatin and histone marks observed after decidualization correspond to regulatory elements that become more or less active, resulting in correlated gene expression changes of the nearby genes.

**Figure 2.**
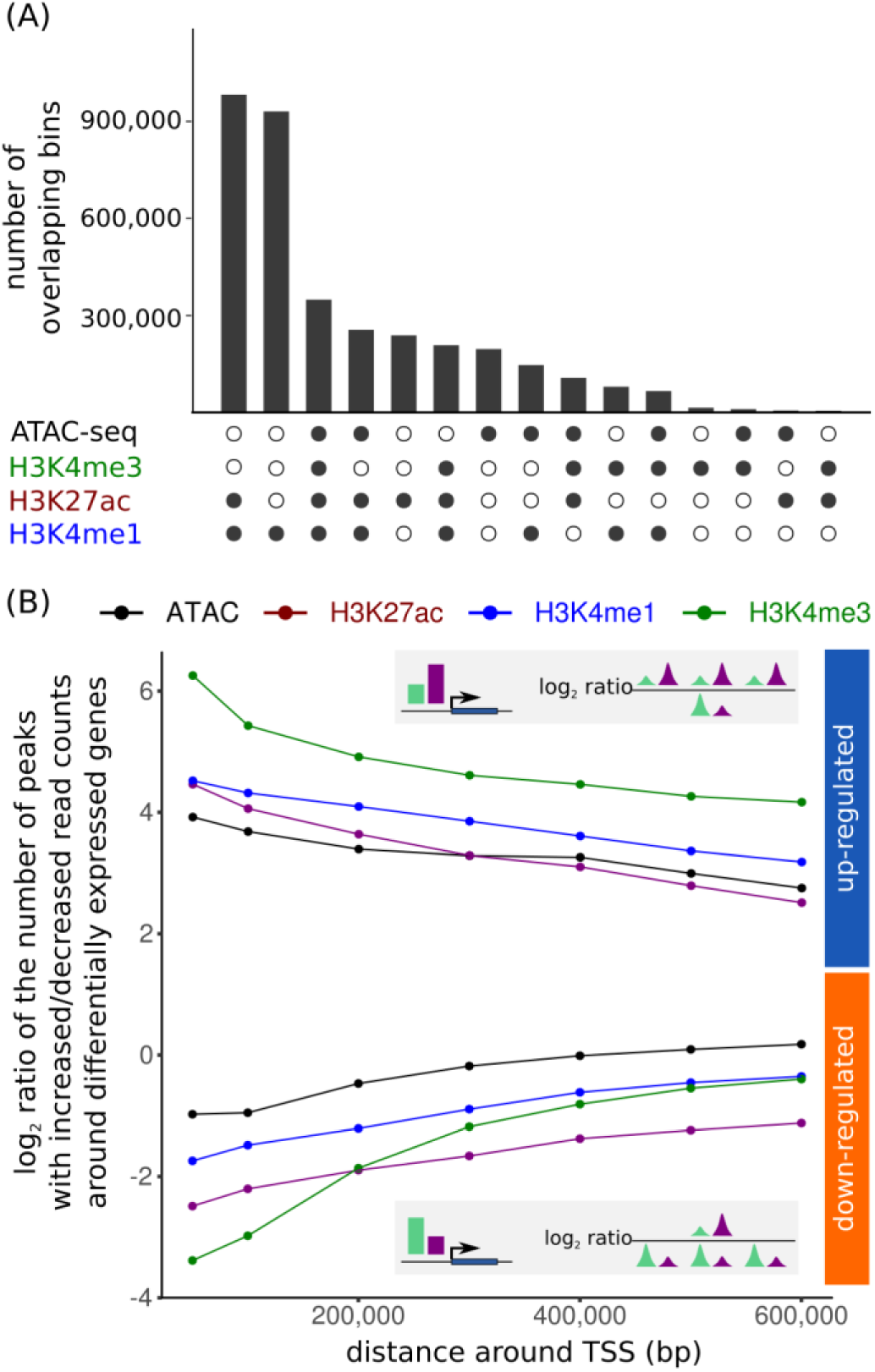
Characteristics of ATAC-seq and histone modification maps in response to decidualization. (a) Plot showing the overlap between the different histone modifications and ATAC-seq maps. Peaks were assigned to 100 bp bins to avoid ambiguity in overlap due to different peak borders. (b) Each data point shows the ratio between the number of increased/decreased differential peaks nearby genes that increase expression after decidualization (blue, positive log ratios; upper half of the figure) or decrease expression after decidualization (orange, negative log ratios; lower half of the figure). Genes that were more highly expressed in decidualized cells were flanked by a higher number of ChIP-seq and ATAC-seq peaks that displayed increased read counts in decidualized samples compared to peaks that displayed decreased read counts (top inset). Genes that were down-regulated in decidualized cells showed the opposite trend (bottom inset). All enrichments were highly significant (p < 10^−25^).

Previous work identified transcription factors that play critical roles in decidualized stromal cells^27-31^. Several of the DNA binding motifs that were enriched in peaks with increased or decreased read counts in our data correspond to transcription factors previously implicated in decidualization (Figure 3a), such as CEBP (CAAT-enhancer-binding protein)^32^, PGR (progesterone receptor)^27^, which shares the same motif with ARE (androgen response element) and GRE (glucocorticoid receptor), FOSL2^27^, which shares the same motif with Fra1, Atf3 and BATF, and TEA domain transcription factors^20,33^. Whereas CEBP and PGR were exclusively enriched in peaks with increased read counts in decidualized cells, the FOSL2 motif was present in peaks that both changed positively and negatively in decidualized cells.

**Figure 3.**
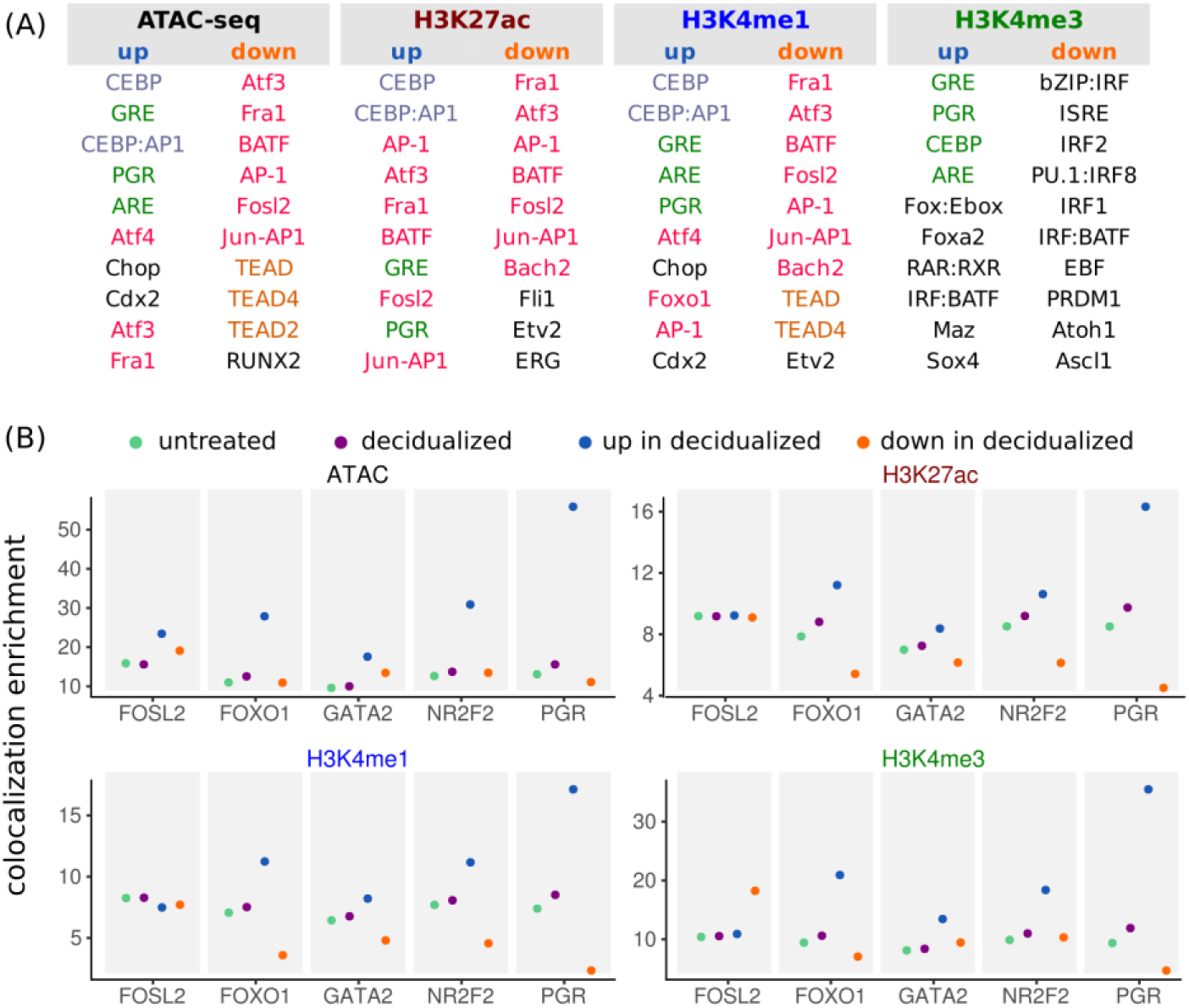
Differential histone modification and ATAC-seq peaks are enriched for transcription factors with roles in decidualization. (a) DNA binding motifs of transcription factors relevant in decidualization are enriched in peaks that change following decidualization treatment. Motifs are color coded by similarity. (b) Colocalization of PGR, FOSL2, FOXO1, GATA2 and NR2F2 with ATAC-seq and ChIP-seq peaks. Transcription factors co-occur with ATAC-seq and ChIP-seq peaks more often compared to co-occurrence with random peaks in both untreated (green) and decidualized (purple) cells. Enrichment of the co-occurrences of PGR, FOXO1, GATA2 and NR2F2 are higher when co-occurring with peaks that have increased read counts (navy blue) and lower with peaks that have decreased read counts (orange) in decidualized compared to untreated cells. Enrichment of co-occurrences with peak sets was calculated as the fold-difference between the number of transcription factor peaks overlapping with ATAC-seq/ChIP-seq peaks and with a random set of peaks (see Methods).

To better understand the role of these transcription factors in decidualization, we obtained publicly available ChIP-seq data for PGR^27^, FOSL2^27^, FOXO1^28^, NR2F2^29^ and GATA2^30^ from endometrial biopsies and analyzed the colocalization of their binding locations with the putative regulatory elements identified by ATAC-seq and ChIP-seq identified in our study (Figure 3b). With the exception of FOSL2, the colocalization enrichments of PGR, FOXO1, GATA2, NR2F2 with ATAC-seq and ChIP-seq peaks were higher (9- to 16-fold) among peaks that were increased in decidualized cells (more open chromatin or increased histone modification levels) compared to all peaks (7.5 to 12.8-fold) and to peaks that decreased in decidualized cells (2- to 5-fold). This observation supports the notion that these transcription factors are involved in regulation of decidualization^27-30,34^. Although FOSL2 has been reported as a positive co-regulator of PGR^27^, the presence of FOSL2 motifs in peaks that both increased and decreased in decidualized cells (Figure 3a) and the lack of difference in the colocalization enrichment between these two sets of peaks (Figure 3b) suggests that FOSL2 may have a dual role in decidualization.

Taken together, our results support a model of decidualization that involves changes in the regulatory landscape during the differentiation of MSCs into DSCs, including alterations in chromatin accessibility and in the activation levels of distant regulatory elements, accompanied by the differential binding of key transcription factors, resulting in increases or decreases in gene expression.

### Chromatin interactions aid in the identification of target genes of distal regulatory elements

As shown in Figure 2, the surrounding regions of differentially expressed genes were enriched for differential ChIP-seq and ATAC-seq peaks that changed in the same direction as the genes in decidualized samples. Accordingly, when we paired differential peaks with the nearest expressed gene as its putative gene target, we observed that these pairs were more likely to have matching directions of change (i.e. both the peak and the gene have increased or decreased read counts in decidualized samples) than non-matching directions when compared with pairs that were assigned randomly (Figure 4A).

**Figure 4.**
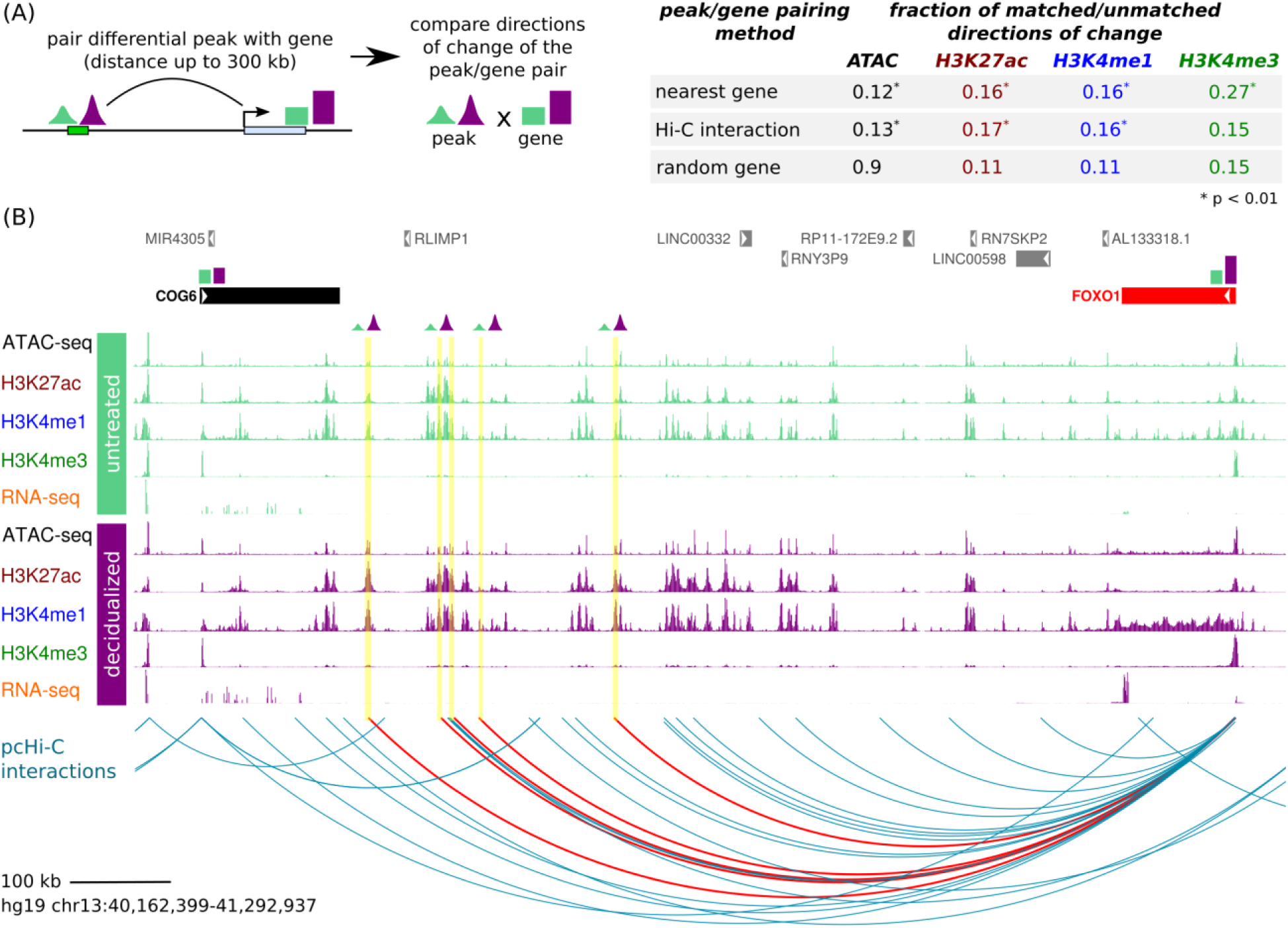
pcHi-C connects predicted regulatory elements to their putative target genes. (A) Randomly assigning a gene to a peak (see Materials and Methods) resulted in fewer peaks that matched the direction of change with that of differentially expressed genes than when using pcHi-C interactions or the nearest gene to pair peaks to genes. (B) The *FOXO1* gene is more highly expressed in decidualized samples and its promoter physically interacts (red arcs) with distal regulatory elements (yellow highlights) that show increased activation in decidualized samples. The nearest expressed gene to these differential peaks is *COG6*.

In many cases, however, the target gene for a regulatory element is not the nearest gene^35^ and therefore, information about distal chromatin interactions can be useful in prioritizing candidate gene targets of variants identified in GWAS. To this end, we generated a promoter capture Hi-C map (pcHi-C) of a decidualized cell line, thus enriching for the identification of long-range chromatin interactions between promoters and distant regulatory elements^36-38^. We identified a total of 161,337 interactions, of which 53,211 were between promoters and distal regions of accessible chromatin assayed by ATAC-seq and ChIP-seq, suggestive of their regulatory role. We used the significant interactions between promoters and distal regions that we identified to pair differential peaks with putative target genes. As shown in Figure 4A, using pcHi-C interactions as a pairing method resulted in enhanced identification of differential peak/differential target gene pairs that have matching directions of change compared to random assignment of gene-target pairs.

Whereas assigning peaks to the nearest expressed gene also led to enhanced assignment of differential peaks to target genes with matching directions of change (Figure 4A), pcHi-C was helpful in identifying less obvious target genes, as show in Figure 4B. In this example, several pcHi-C interactions link distal regulatory elements up to 847 kb away that became more active in decidualized cells to the promoter of a gene (*FOXO1*) that was up-regulated in decidualized cells and is known to be involved in decidualization^31^. The nearest expressed gene method assigned those differential peaks to *COG6*, a gene that does not change expression in decidualized samples and is therefore a less likely target.

In conclusion, by combining pcHi-C interactions with the epigenome maps and transcriptome data we were able to identify genes and putative regulatory elements that respond to, or regulate, the decidualization process. We next used these comprehensive functional genomic maps and datasets to fine map GWAS loci for gestational duration and identify new candidate genes with a potential role in PTB.

### The heritability of gestational duration is enriched for functional annotations in decidual stromal cells

To identify candidate genes that may play a role in gestational duration and PTB, we used summary data from a GWAS of gestational duration based on a meta-analysis of a 23andMe GWAS (N=42,121)^5^ and the results from six European data sets (N=14,263). A detailed description of the GWAS is in Supplementary File S3 and Supplementary Figures S2 and S3. After filtering for SNPs that are present in the 1000 Genomes Project data and minor allele frequency (MAF) > 0.01, we identified SNPs at six autosomal loci, defined as approximately independent blocks by LDetect^39^, that were associated with gestational duration at genome-wide significance of p < 5×10^−8^ (Supplementary Table S1). We then created a computational pipeline to assess enrichment of GWAS signals in functional annotations that we generated in untreated (MSCs) and decidualized (DSCs) stromal cells to fine map GWAS loci and discover candidate causal genes, and to potentially provide support for additional loci that did not reach genome-wide significance in the GWAS (Figure 5).

**Figure 5.**
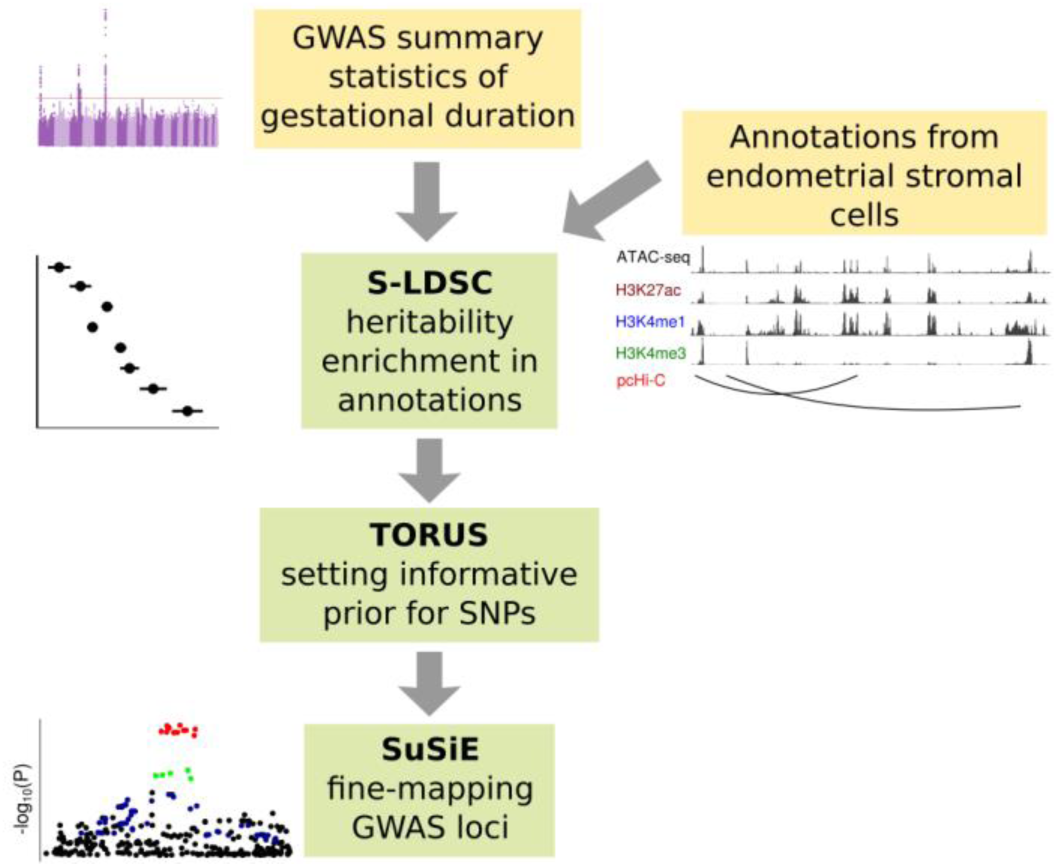
Computational pipeline of our GWAS analysis. Yellow boxes (input data): GWAS summary statistics and functional annotations from endometrial stromal cells (in both untreated and decidualized cells). Green boxes: steps of computational processing.

We first used Stratified-LD score regression (S-LDSC)^40^ to assess enrichment of GWAS signals in functional annotations in endometrial stromal cells. S-LDSC is a commonly used tool for estimating the proportion of heritability of complex phenotypes that is explained by variants in certain functional annotations. The heritability enrichment is defined by the proportion of heritability explained by annotations divided by the expected proportion, which is the percent of SNPs genome-wide that are in these functional annotations. Importantly, to account for possible systematic bias in this analysis, i.e. SNPs within annotations of interest may differ from background SNPs in systematic ways such as their LD structure and epigenomic properties, we included a range of baseline annotations (default S-LDSC setting), including LD-related annotations, DNase hypersensitivity, enhancer annotation, H3K27ac, H3K4me1 and other histone marks (the union across cell types). Thus, if an annotation is shared by many cell types, it would not show enrichment in S-LDSC analysis (see Methods).

Using S-LDSC, we found 5- to 10-fold enrichments of GWAS heritability for gestational duration in our functional annotations compared to the baseline model of S-LDSC (Figure 6). The enrichment of enhancer marks H3K27ac and H3K4me1 were higher in decidualized than in untreated cells, but the opposite pattern was observed for the promoter mark H3K4me3, which was more enriched in untreated (MSCs) than in decidualized (DSCs) cells. These findings are consistent with previous observations that enhancers are often more dynamic and condition- or tissue-specific than promoters^10^. We observed weaker heritability enrichments of open chromatin regions defined by ATAC-seq and of interaction regions in pcHi-C. However, because we performed joint analysis of all annotations together, the enrichment of one annotation (e.g. ATAC-seq peaks) will be reduced if the enrichment is partially explained by other, overlapping annotations (e.g. H3K27ac). Although the promoter mark H3K4me3 in untreated cells showed the highest enrichment, the annotations that contributed most to the heritability of gestational duration were enhancers (Figure 6), due to the much larger number of enhancer histone marks than promoters in the genome. Our results thus highlight the importance of functional annotations in endometrial stromal cells at GWAS loci for gestational duration.

**Figure 6.**
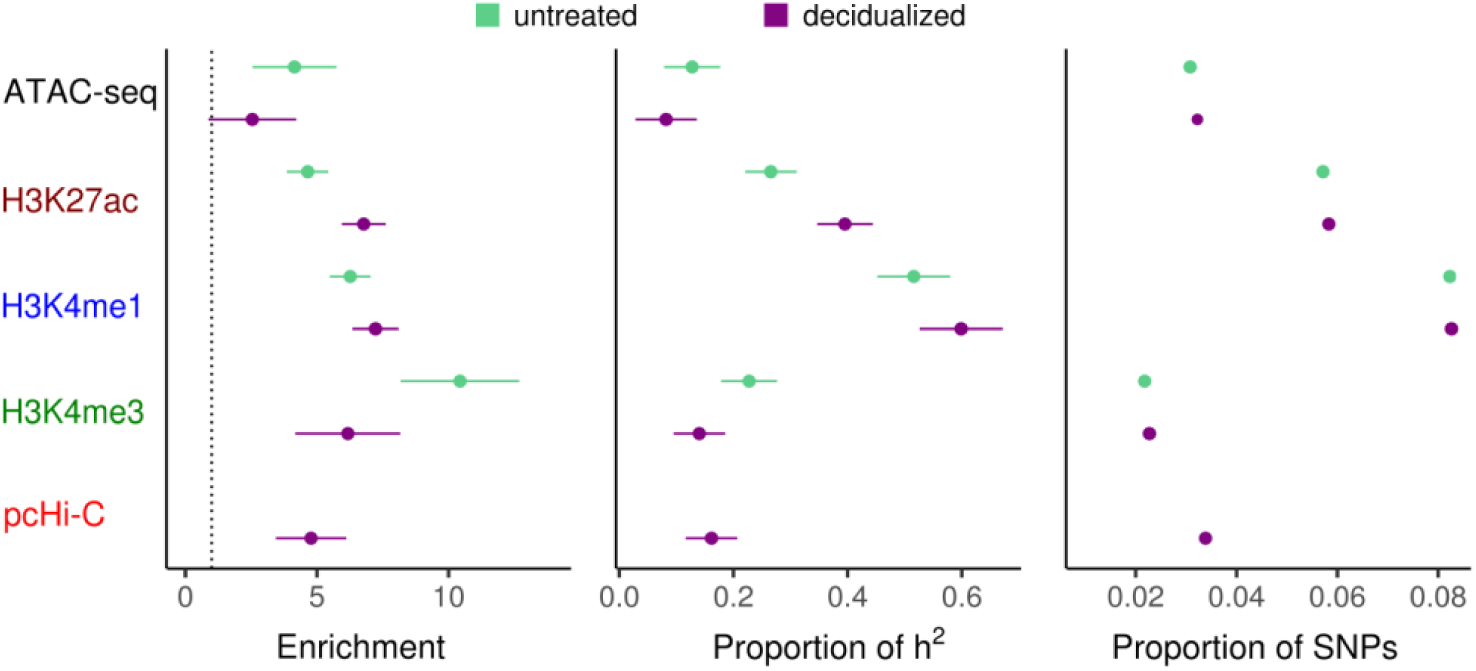
Stratified LDSC heritability analysis of GWAS of gestational duration using functional annotations. Left panel: fold enrichment of heritability in each annotation. Dashed line shows values at 1, i.e. no enrichment. Center: proportion of heritability explained by each annotation. Right: proportion of SNPs across the genome that fall within an annotation. For each annotation, enrichment (left) is the ratio of h^2^ proportion (center) divided by the SNP proportion (right). Error bars represent 95% confidence intervals.

### Integrated analysis of GWAS and decidual cell functional annotations improves fine-mapping of causal variants of gestational duration and identifies putative target genes

We next developed a computational procedure to integrate the decidua stromal cell functional maps with genetic maps of reproductive traits (Figure 5). We posited that integrating functional maps in these pregnancy-relevant cells and leveraging statistical methods to fine-map associations would result in 1) identifying candidate causal variants in each associated locus, 2) linking those variants to their target genes, and 3) discovering additional loci and genes associated with gestational duration.

We first leveraged the enrichments of DSC annotations to create Bayesian prior probabilities for a variant being causal. Using prior probabilities informed by functional annotations of SNPs could increase the accuracy of fine-mapping, as shown in recent studies^8,41^. Based on the results of S-LDSC, we chose H3K27ac, H3K4me1 and pcHi-C interactions from the decidualized cells, and H3K4me3 from untreated cells, as functional genomic annotations to create informative priors using TORUS^42^. To assign a prior to each SNP, TORUS uses genome-wide summary statistics of GWAS and the functional annotations to assess how informative each annotation is in predicting causal variants (Supplementary Table 2). SNPs associated with functional annotations are generally assigned higher prior probabilities. Additionally, TORUS computes statistical evidence at the level of genomic blocks, defined as the probability that a block (determined by LD) contains at least one causal SNP. Without including any histone marks or chromatin accessibility annotations, TORUS implicated six autosomal blocks in the genome at FDR < 0.05, including five of the six genome-wide significant autosomal loci identified in the GWAS (p < 5×10^−8^). One locus on chromosome 3 had an FDR = 0.11, and was therefore not identified by TORUS and one locus on chromosome 9 that was not identified in the GWAS was implicated by TORUS (Supplementary File S4). By including the functional genomic annotations from endometrial stromal cells, the number of high confidence blocks increased to ten, including all six that were significant in the gestational duration GWAS and four that were not significant in the GWAS (Supplementary File S4).

**Table 2.**
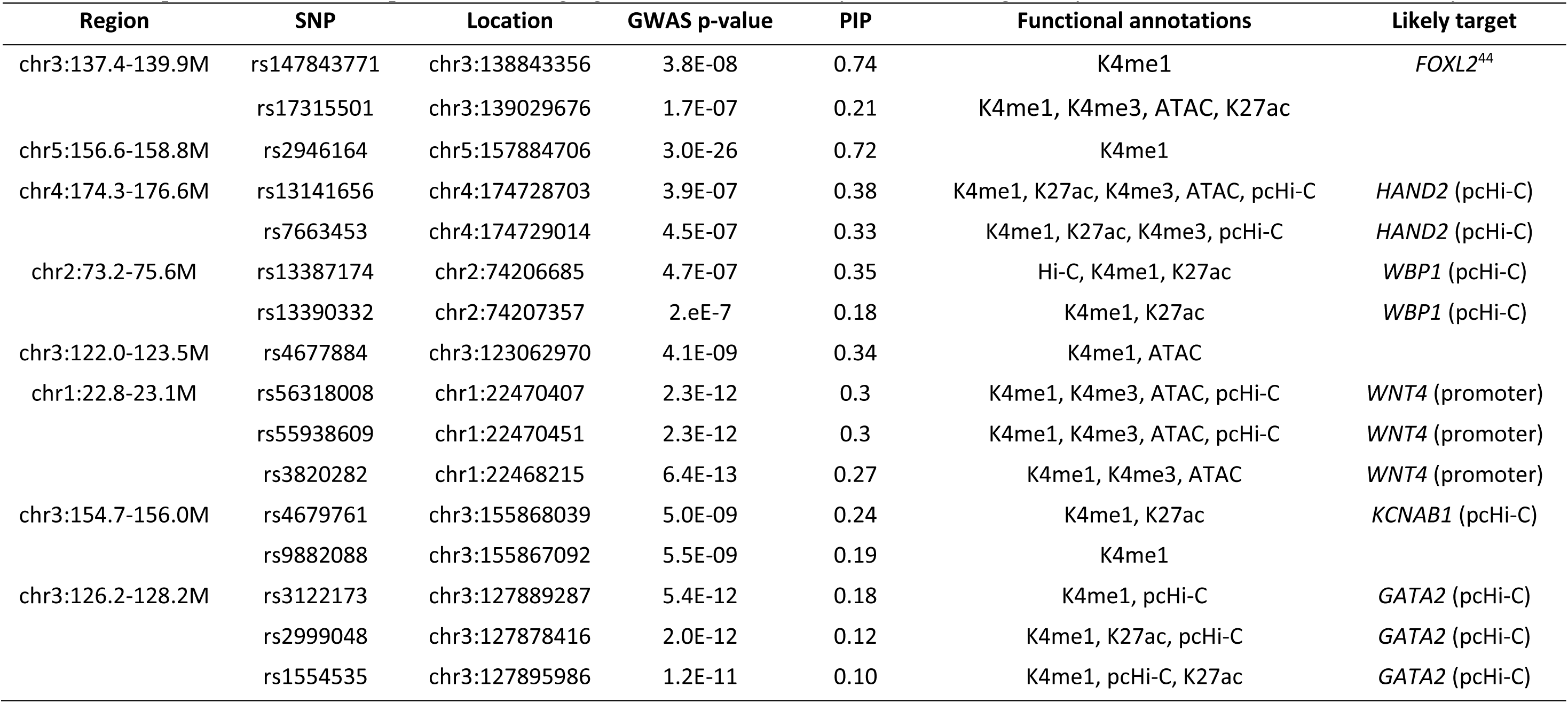
Most probable SNPs identified from computational fine-mapping of regions associated with gestational duration. Functional annotations are based on data from endometrial stromal cells. We list an annotation if the SNP is located in a sequence with that annotation in either untreated or decidualized condition. We list the pcHi-C annotation if the SNP is within 1 kb of a region involved in a pcHi-C interaction. We call a gene the target of a SNP if (1) the SNP is located in the promoter (< 1 kb of TSS) of that gene; or (2) the promoter of that gene has a pcHi-C interaction with a region within 1 kb of the SNP. In the case of rs147843771 at the *FOXL2* locus, the target was defined by literature evidence^44^. The number of credible SNPs at each region are shown in Figure 7B. *FOXL2*: Forkhead box L2, *GATA2*: GATA-binding protein 2, *HAND2*: Heart and Neural Crest Derivatives Expressed 2, *KCNAB1*: potassium voltage-gated channel subfamily A member regulatory beta subunit 1, *WNT4*: Wnt family member 4.

We next performed computational fine mapping on these ten blocks, with the informative priors learned by TORUS, using SuSiE^43^. Conceptually, SuSiE is a Bayesian version of the stepwise regression analysis commonly used in GWAS (i.e. conditioning on one variant, and testing if there is any remaining signal in a region). SuSiE accounts for the uncertainty of causal variants in each step, and reports the results in the form of posterior including probabilities (PIPs). The PIP of a variant ranges from 0 to 1, with 1 indicating full confidence that the SNP is a causal variant. If a region contains a single causal variant, the PIPs of all SNPs in the region should approximately sum to 1.

Including the priors defined by TORUS using DSC functional annotations significantly improved fine-mapping (Figure 7A, Supplementary Table S2 and Supplementary File S5. For example, only one SNP reached PIP > 0.3 across all 10 blocks using the default setting under SuSiE (uniform prior, treating all SNPs in a block equally). This reflects the general uncertainty of pinpointing causal variants due to LD: e.g., a strong GWAS SNP in close LD with 9 other SNPs would have PIP about 0.1. By using the annotation-informed priors, 8 SNPs in six different blocks reached PIP > 0.3 (Figure 7A). In some blocks, we were able to fine-map a single high-confidence SNP, e.g. the *FOXL2* locus on chromosome 3, while in other blocks, we had considerable uncertainty of the causal variants, as shown by large credible sets, i.e. the minimum set of SNPs to include the causal SNP with 95% probability (Figure 7B). Table 2 summarizes the most probable causal variants in eight blocks (fine-mapping in the remaining two blocks produced large credible sets with no high-PIP SNPs) as well as their likely target genes based on promoter assignment or chromatin interactions from pcHi-C. We note that our results of the *WNT4* locus identified rs3820282 as the likely causal variant. This is consistent with our previous results demonstrating experimentally that the T allele of this SNP disrupts the binding of estrogen receptor 1 (ESR1)^5^. This SNP was among the 3 most likely SNPs in our fine-mapping study, with a PIP of 0.27 (Table 2).

**Figure 7.**
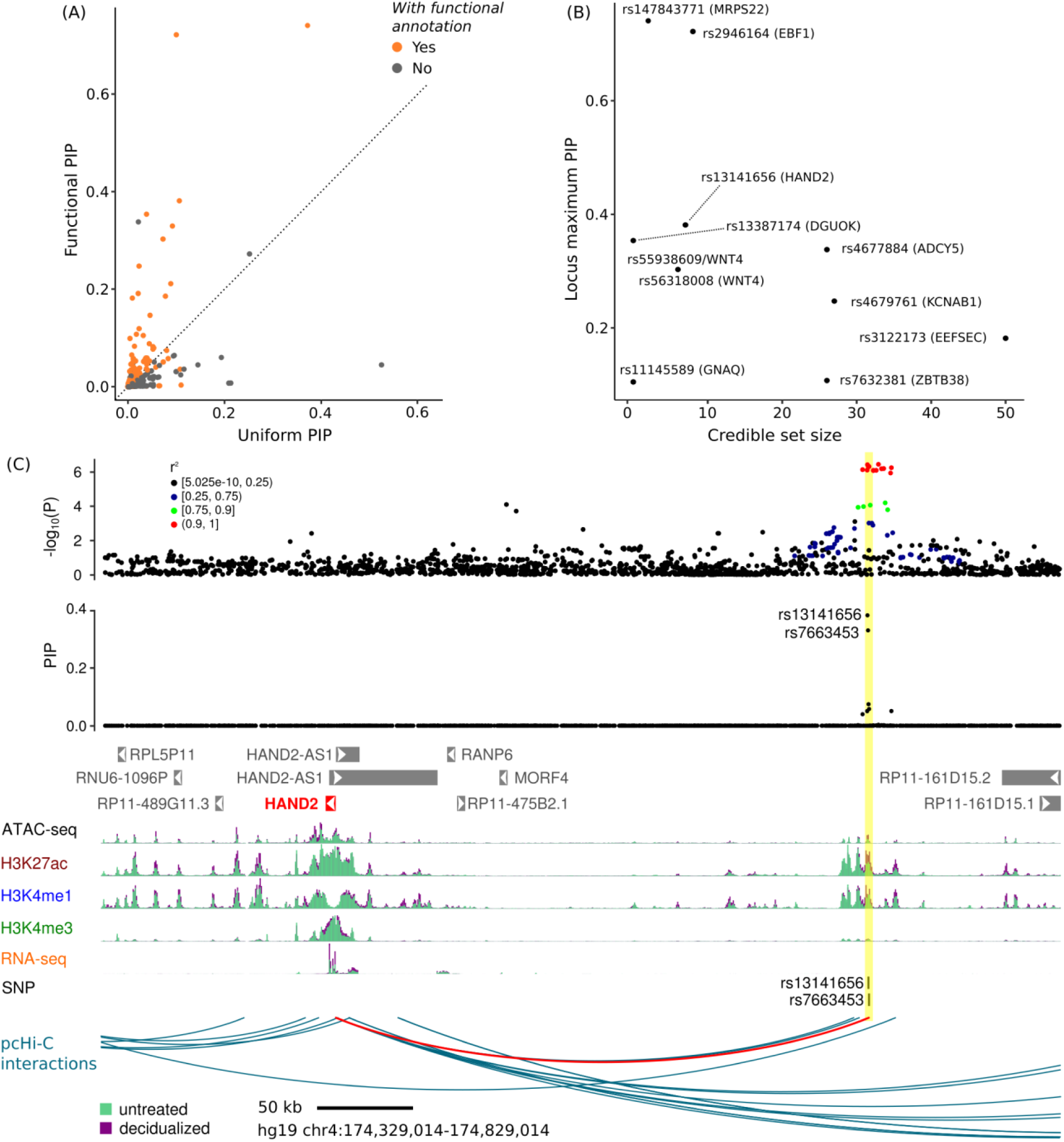
Fine-mapping GWAS loci of gestational duration. (A) PIPs of SNPs using uniform vs. functional priors in SuSiE (each dot is a SNP). The functional prior of a SNP is based on SNP annotations and is estimated using TORUS. (B) Summary of fine-mapping statistics of all 10 regions. X-axis: the size (number of SNPs) of credible set. Y-axis: the maximum PIP in a region. We label each region by its top SNP (by PIP) and the likely causal gene, according to Table 2 or the nearest gene of the top SNP. (C) Likely causal variants near *HAND2* and their functional annotations. The upper panel shows the significance of SNP association in the GWAS and the middle panel shows fine-mapping results (PIPs) in the region. The vertical yellow bar highlights the two SNPs with high PIPs. These SNPs are located in a region annotated with ATAC-seq, H3K27ac, H3K4me1 and H3K4me3 peaks (bottom). This putative enhancer also had increased ATAC-seq, H3K27ac and H3K4me1 levels in decidualized samples and interacts with the *HAND2* promoter (red arc).

We highlight the results from two regions. In the first, two adjacent SNPs (311 bp apart), rs13141656 and rs7663453, on chromosome 4q34 did not reach genome-wide significance in the GWAS (p = 3.9 ×10^−7^ and 4.5 ×10^−7^, respectively). After using functional annotations in decidua-derived stromal cells, the block containing these SNPs was highly significant (TORUS q-value = 0.02), suggesting the presence of at least one causal variant in this block. The two SNPs together explained most of the PIP signal in the block (PIP 0.38 and 0.33, respectively, Table 2). The two SNPs are located in a region of open chromatin in endometrial stromal cells, with enhancer activity marked by both H3K27ac and H3K4me1 (Figure 7C). Only 9 of the 129 tissues from the Epigenome Roadmap^11^ also had H3K27ac, H3K4me1 or H3K4me3 peaks spanning the rs13141656 locus and only 2 spanning the rs7663453 locus. In addition, this putative enhancer is bound by multiple transcription factors, including GATA2, FOXO1, NR2F2 and PGR, based on ChIP-seq data. The only physical interaction of this enhancer in the pcHi-C data in decidualized stromal cells is with the promoter of the *HAND2* gene, located 277 kb away (Figure 7C). Summing over the PIPs of all SNPs whose nearby sequences interact with *HAND2* via chromatin looping gives an even higher probability, 0.89, suggesting that *HAND2* is very likely to be the causal gene in this region (Supplementary Table S3). HAND2 is an important transcription factor that mediates the effect of progesterone on uterine epithelium^45^. Thus, in this example we identified a novel locus, the likely causal variant(s), the enhancers they act on, and an outstanding candidate gene for gestational duration and PTB.

The second example focuses on the locus showing a strong GWAS association with gestational duration on chromosome 3q21. The lead SNP, rs144609957 (GWAS p = 4 ×10^−13^), is located upstream of the *EEFSEC* (Eukaryotic Elongation factor, Selenocysteine-TRNA Specific) gene. There is considerable uncertainty of the causal variants in this region, with 50 SNPs in the credible set and the lead SNP explaining only a small fraction of signal (PIP = 0.02). Among all 12 SNPs with PIP > 0.01, 11 have functional annotations, most commonly H3K4me1 and pcHi-C interactions. Interestingly, for nine SNPs (first 3 shown in Table 2), the sequences in which they are located physically interact with the promoter of *GATA2* in the pcHi-C data, but not with any other promoters in the region (Supplementary Figure S4). The PIPs of all SNPs in the genomic regions that likely target *GATA2* through chromatin looping sum to 0.68 (Supplementary Table S4). Thus, despite uncertainty of causal variants in this region, our results implicate *GATA2* as a candidate causal gene in endometrial stromal cells. GATA2 is a master regulator of embryonic development and differentiation of tissue-forming stem cells^46^. As support for the possible role of GATA2 in pregnancy, *GATA2* deficient mice show defects in embryo implantation and endometrial decidualization^34^, making this another excellent candidate causal gene for gestational duration and PTB.

## Discussion

The molecular processes that signal the onset of parturition in human pregnancies, and how perturbation of those processes result in PTB, are largely unknown. Yet, understanding such processes would reveal important insights into the potential causes of adverse pregnancy outcomes, including spontaneous labor prior to 37 weeks’ gestation, and potentially lead to the identification of biomarkers and therapeutic targets for PTB. Unbiased genome-wide association studies do not require prior knowledge of molecular processes underlying disease phenotypes and have the potential to identify novel genes and pathways contributing to common diseases. However, the significant heterogeneity of most common diseases and small effects of most common disease-associated variants lead to the requirement for very large sample sizes (in the tens to hundreds of thousands of cases) to discover more than a handful of associated loci that meet stringent criteria for genome-wide significance. To address this limitation and provide orthogonal evidence for assessment of associations, we characterized the transcriptional and chromatin landscapes in decidua-derived stromal cells and integrated those functional annotations with a GWAS of gestational duration to discover novel loci and genes. The primary motivation for these studies was the striking paucity of genomic and epigenomic functional annotations in pregnancy-relevant primary cells among those studied by large consortia^9-11^. Here, we filled a significant gap by providing comprehensive maps in untreated and decidualized stromal cells, and used these maps for annotating GWAS of pregnancy-related traits.

We chose to focus these studies on endometrial stromal cells because of their central importance in both the establishment and maintenance of pregnancy, as well as their intimate juxtaposition to fetal trophoblast cells throughout pregnancy. Of particular relevance are the roles that decidualized stromal cells play in regulating trophoblast invasion, modulating maternal immune and inflammatory responses at the maternal-fetal interface, and controlling remodeling of the endometrium^47^. In fact, defects in all of these processes have been considered a contributing factor to pregnancy disorders^47,48^. Moreover, we showed that the SNPs in regions with endometrial stromal cell functional annotations explained more of the heritability of gestational duration compared to just using baseline annotations. Among all annotations, enhancer marks H3K4me1 (in both decidualized and untreated stromal cells) and H3K27ac (in decidualized cells) were 8- to 10-fold enriched at GWAS loci after adjusting for the general annotations, and accounted for 50-70% of the GWAS heritability. The lack of complete independence between these marks makes it difficult to delineate their individual effects but nonetheless highlights the importance of enhancers and of gene regulation in endometrial stromal cells in modulating the effects of GWAS variants on gestational duration. This is consistent with both the known tissue-specific roles of enhancers and the observation that over 90% of GWAS loci reside outside of the coding portion of the genome and are enriched in regions of open chromatin and enhancers^12,40^.

Integrating transcriptional and chromatin annotations of gene regulation from MSCs and DSCs improved our ability to discover novel GWAS loci and identify likely causal SNPs and genes associated with gestational duration. We illustrate how our integrated platform identified a novel causal locus and candidate gene (*HAND2*) associated with gestational duration, as well as refined the annotation of loci that had been previously identified. Our data suggest that in endometrial stromal cells *GATA2* is likely the target gene of enhancers harboring SNPs associated with gestational duration. This does not exclude the possibility that the nearest gene to the associated SNPs, *EEFSEC*, may be a target gene in other cell types.

Both of these examples highlight transcription factors that are essential for endometrial development or decidualization. The fact that neither *GATA2* nor *HAND2* were identified as potential candidate genes in previous GWASs of gestational duration or PTB supports our approach and the importance of using functional annotations from cell types relevant to pregnancy to fine map and identify candidate genes for the pregnancy-related traits. Overall, the integrated analyses performed in this study resulted in the identification of both novel GWAS loci and novel candidate genes for gestational duration, as well as maps of the regulatory architecture of these cells and their response to decidualization.

However, there are some limitations. In particular, we focused on only one cell type, albeit one that plays a central role in pregnancy, and only one exposure (hormonal induction of decidualization). Future studies that include fetal cells from the placenta and uterine or cervical myometrial cells could reveal additional processes that contribute to gestational duration and PTB, such as those related to fetal signaling and the regulation of labor, respectively. Inclusion of additional exposures, such as trophoblast conditioned media^49^, may further reveal processes that are pregnancy-specific. Second, to maximize power we focused on a GWAS of gestational duration and not PTB *per se*. While previous GWAS have shown that all PTB loci were among the gestational age loci^5^, we realize that some of the loci that we identified could be related to normal variation in gestational duration and not specifically to PTB. Nonetheless, our findings contribute to our understanding of potential mechanisms underlying the timing of human gestation, about which we still know little. Lastly, although our ChIP-seq results revealed an association between GATA2 binding and decidualization, confirming the role of this transcription factor in decidual cell biology^50,51^, and studies in murines support its role in endometrial processes^34^, we do not yet have direct evidence showing that perturbations in the expression of *GATA2*, or any of the other target genes identified, influence the timing of parturition in humans. Future studies will be needed to directly implicate the expression of these genes in gestational duration or PTB.

In summary, our study highlights the importance of generating functional annotations in pregnancy-relevant cell types to inform GWASs of pregnancy-associated conditions. Our results suggest that the expression of two transcription factors, *GATA2* and *HAND2*, in endometrial stromal cells may regulate transcriptional programs that influence the timing of parturition in humans, which could lead to the identification of biomarkers of or therapeutic targets for PTB.

## Materials and methods

### Sample collections

Placentas were collected from three women (≥18 years old) who delivered at term (≥37 weeks) following spontaneous labor; all were vaginal deliveries of singleton pregnancies. Within 1 hour of delivery, 5 x 5 cm pieces of the membranes were sampled from a distant location of the rupture site. Pieces were placed in DMEM-HAMS F12 media containing 10% FBS and 1% pen/strep. Samples were kept at 4°C and processed within 24 hours of tissue collection. This study was approved by the Institutional Review Boards at the University of Chicago, Northwestern University and Duke University Medical School.

### Isolation of mesenchymal stromal cells from human placental membranes

Third trimester placental tissue was enzymatically digested by a modification of previously described methods^52,53^. Decidua tissue was gently scraped from chorion, and tissue was enzymatically digested in a solution (1X HBSS, 20mM HEPES, 30mM sodium bicarbonate, 1% BSA fraction V) containing collagenase type IV (200 U/mL; Sigma *C-5138*); hyaluronidase Type IS (1mg/mL; *Sigma H-3506)* and DNase type IV (0.45 KU/mL, *Sigma D-5025)* at 37°C, until a single cell suspension was obtained (usually 3 rounds of 30 min digestion using fresh digestion media each round). Epithelial cells were removed by filtering through a 75uM nylon membrane and RPMI (Sigma) containing 10% FBS was added for enzyme inactivation. Dissociated cells were collected by centrifugation at 400g for 10 minutes and washed in RPMI/10%FBS. Erythrocytes were removed by cell pellet incubation with 1X Red Blood Cell Lysis Buffer (Sigma) for 2.5 minutes at room temperature. The resulting cells were counted and resuspended in seeding media (1x phenol red-free high glucose DMEM (GIBCO) supplemented with 10% FBS (Thermofisher), 2 mM L-Glutamine (Life Technologies), 1 mM Sodium Pyruvate (Fisher), 1x Insulin-Transferrin-Selenium (ITS; Thermofisher), 1% pen-strep and 1x anti-anti (Thermofisher). Dissociated cells were plated into a T75 flask and incubated at 37°C and 5% CO2 for 15-30 min (enrichment by attachment). The supernatant was carefully removed and loosely attached cells were discarded. Plates were allowed to grow in fresh media containing 10% charcoal stripped FBS (CS-FBS) and 1x anti-anti until the plate was 80% confluent. The anti-anti was removed from the culture media after 2 weeks of culture. We obtained >99% vimentin-positive cells after 3 passages (Supplementary Figure S1). Cells were expanded, harvested in 0.05% trypsin and cryopreserved in 10% dimethylsulfoxide culture media for subsequent use. Each cell line was defined as coming from a different sample collection (different pregnancy).

### Decidualization of mesenchymal stromal cells in vitro

Cells were plated and grown for 2 days in cell culture media (1x phenol red-free high glucose DMEM, 10% CS-FBS, 2 mM L-Glutamine, 1 mM Sodium Pyruvate and 1x ITS). After 2 days, cells were treated either with control media (1x phenol red-free high glucose DMEM, 2% CS-FBS, 2mM L-Glutamine) or decidualization media (1x phenol red-free high glucose DMEM, 2% CS-FBS, 2mM % L-Glutamine, 0.5mM 8-Br-cAMP, 1uM MPA) for 48 hours. Cells were incubated at 37°C, 5% CO2 and harvested for ATAC-seq, ChIP-seq and RNA-seq. Experiment with 3 different cell lines was performed in triplicate and repeated 3 times. Prolactin (PRL) and IGFBP1 mRNA were assessed by quantitative real-time PCR before each downstream assay was performed.

### RNA-seq

Total RNA was extracted from approximately 1 million cells using AllPrep DNA/RNA Kit (Qiagen) according to manufacturer’s instructions. RNA quality (RNA integrity number - RIN) and concentration was assessed by Bioanalyzer 2100 (Agilent technology). RNA-seq libraries were generated by TruSeq stranded total RNA library prep kit (Illumina) and TruSeq RNA CD Index Plate.

### ChIP-seq

For ChIP experiments, cells were crosslinked by adding to the media 37% formaldehyde to a final concentration of 1%, gently mixed, incubated for 10 minutes, and quenched for 5 minutes with 2.5M glycine for a final of 0.125M per plate. Cells were washed using cold 1X PBS, and scraped in 15mL cold Farnham lysis buffer plus protease inhibitor (Roche, 11836145001), and cell pellets were flash frozen and kept at −80C. Thawed pellets were resuspended in RIPA buffer on ice, aliquoted into 20 million cell per tube, and sonicated by Bioruptor (three 15-minute rounds of 30 seconds ON, 30 seconds). ChIP was performed on 10 million cells using antibodies to H3K27ac, H3K4me3, and H3K4me1 histone marks (ab4729/lot# GR274237, ab8580/lot# GR273043 and ab8895/lot# GR262515, respectively). M-280 sheep anti-rabbit IgG Dynabeads (Invitrogen, 11203D) was used for chromatin immunoprecipitation. DNA was purified using Qiagen MinElute PCR Purification Kit, quantified by Qubit, and prepared for sequencing using the Kapa Hyper Prep Kit. All libraries were pooled to 10nM per sample prior to sequencing.

### ATAC-seq

Approximately 50,000 cells were harvested and used for ATAC-seq library preparation as described in the Fast-ATAC protocol^54^. ATAC-seq libraries where uniquely indexed with Nextera PCR Primers and amplified with 9 to 12 cycles of PCR amplification. Amplified DNA fragments were purified with 0.8:1 ratio of Agencourt AMPure XP (Beckman Coulter) to sample. Libraries were quantified by Qubit, and size distribution was inspected by Bioanalyzer (Agilent Genomic DNA chip, Agilent Technologies). All libraries were pooled to 10nM per sample prior to sequencing.

### Promoter capture Hi-C

In situ Hi-C was performed as described previously^55^. Briefly, 5 million decidualized cells were treated with formaldehyde 1% to cross-link interacting DNA loci. Cross-linked chromatin was treated with lysed and digested with MboI endonuclease (New England Biolabs). Subsequently, the restriction fragment overhangs were filled in and the DNA ends were marked with biotin-14-dATP (Life Technologies). The biotin-labeled DNA was sheared and pulled down using Dynabeads MyOne Stretavidin T1 beads (Life Technologies, 65602) and prepared for Illumina paired-end sequencing. The *in situ* Hi-C library was amplified directly off of the T1 beads with 9 cycles of PCR, using Illumina primers and protocol (Illumina, 2007). Promoter capture was performed as described previously^38^. The Hi-C library was hybridized to 81,735 biotinylated 120-bp custom RNA oligomers (Custom Array) targeting promoter regions (4 probes/RefSeq transcription start sites). After hybridization, post-capture PCR was performed on the DNA bound to the beads via biotinylated RNA.

### Differential expression

Read counts per gene were calculated with Salmon^56^ version 0.12.0 on transcripts from human Gencode release 19 (ftp://ftp.ebi.ac.uk/pub/databases/gencode/Gencode_human/release_19/gencode.v19.pc_transcripts.fa.gz and ftp://ftp.ebi.ac.uk/pub/databases/gencode/Gencode_human/release_19/gencode.v19.lncRNA_transcripts.fa.gz). Estimated counts were used in exploratory analysis (transformed with DESeq2’s *rlog* function) and in DESeq2^24^ version 1.24.0 to identify differentially expressed genes (adjusted p-value ≤ 0.05 and absolute fold-change of ≥ 1.2). After observing that replicates for each cell lines clustered together, we pooled reads for each cell line, combining 3 decidualization experiments in each sample. We then performed a paired analysis to obtain genes that were differentially expressed between untreated and decidualized samples.

### Peak calling

ATAC-seq reads were trimmed with cutadapt and aligned with bowtie2^57^ version 2.3.4.1. Reads with mapping quality lower than 10 were discarded. ChIP-seq reads were also aligned with bowtie2. Peaks were called using MACS2^58^ version 2.1.2 with parameters --llocal 20000 --shift - 100 --extsize 200 -q 0.05 for ATAC-seq and default parameters for ChIP-seq. Peaks overlapping coordinates blacklisted by Anshul Kundaje were excluded (http://mitra.stanford.edu/kundaje/akundaje/release/blacklists/hg19-human/).

### Differential ATAC-seq and ChIP-seq peaks

Similarly to RNA-seq, we pooled reads from replicates for each cell line. We called peaks for each of the 6 samples using MACS2 and converted peak coordinates into 100 bp contiguous bins. Bins covered by less than 60% of their extension were excluded. To identify reproducible peaks, we only kept bins that were present in at least 2 out of the 3 cell lines in each condition, allowing for condition-specific peaks. We then merged all adjacent bins, expanding them back into longer peaks. We counted the number of reads in all peaks and in all samples and compared the read counts using DESeq2 (adjusted p-value < 0.05 and absolute fold-change > 1.2).

### Statistical analysis of the frequencies of differential peaks near differentially expressed genes

The p-values in Figure 2B were calculated with a chi-square test of the number of peaks with increased or decreased numbers of reads observed and an expected probability based on the number of peaks in each category for each data set. Bonferroni correction was performed to correct for multiple testing.

### Transcription factor ChIP-seq

ChIP-seq reads were downloaded from NCBI GEO and processed locally. HOMER 4.9^59^ was used to call peaks for the following samples: PGR (GSE94038), NR2F2 (GSE52008), FOSL2 (GSE94038), FOXO1 (GSE94037) and NR2F2 input (GSE52008) and FOXO1, PGR, FOSL2 input (GSE94038).

### Overlap between ATAC-seq and ChIP-seq peaks

Reproducible peaks were converted into 100 bp bins and those with >60% of their extension covered by a peak were retained. Common bins were counted and the number of counts were plotted with UpSetR 1.4.0^60^.

### Motif enrichment

We used HOMER 4.9 to identify DNA binding motifs enriched in peaks with parameters -len 8,10,12 -size 200 -mask.

### Enrichment of overlap between peaks

Enrichment was calculated as the observed number of overlapping peaks divided by the expected number of overlapping peaks using bedtools intersectBed^61^ with a 1 bp minimum. The expected number of overlapping peaks was obtained by averaging 100 random samples of peaks with bedtools shuffle excluding gaps annotated by the UCSC Genome Browser^62^. While shuffling peaks does not account for mapping and other biases that make peak locations non-uniform and may result in over-estimation of enrichment, our results are limited to comparisons between enrichments, which should cancel any biases.

### Hi-C interaction calling

We used HiCUP v0.5.9^63^ to align and filter Hi-C reads. HiCUP used bowtie2 version 2.2.3 to align reads. Unique reads were used as input by CHiCAGO^64^ version 1.2.0 and significant interactions were called with default parameters. We only kept interactions identified by CHiCAGO that were in *cis* and with an end located at least 10 kb from a capture probe.

### Pairing differential peaks with putative target genes

Hi-C: significant interactions identified by CHiCAGO that overlapped an ATAC-seq or ChIP-seq peak and were less than 300 kb away from a promoter were used to assign peak/gene pairs. We chose 300 kb because the mean distance between interacting promoter and other regions was 280 kb (median = 200 kb). Nearest gene: bedtools closest -t first -d was used to find the gene closest to a peak, up to 300 kb away. Random gene: all genes up to 300 kb from a peak were selected and one gene was randomly assigned to each peak. For each of these sets of pairs, we calculated the fraction of peak/gene pairs that had the same direction of change according to differential read count analysis with DESeq2, out of the total number of peak/gene pairs. Only genes expressed at >1 transcript per million across all samples were used in the nearest and random gene assignments.

P-values were calculated with a chi-square test comparing the number of cases in the matched and unmatched categories observed in the random set (average from 200 iterations) and in the two peak/gene pairing methods: nearest gene and pcHi-C interactions.

### Gestational duration GWAS

The GWAS results used in this study was an extension of our previously published results^5^. Like our previous study, we utilized summary results from 23andMe, which were obtained from GWAS of gestational duration in 42,121 mothers of European ancestry. In addition, we performed GWA analyses in 14,263 European mothers from six academic data sets. To increase the power of GWA discovery, we performed meta-analysis between the results from 23andMe and the results from the six data sets. See Supplementary File S3 for a full description of the GWAS.

### GWAS enrichment analysis with S-LDSC

We used stratified LD score regression (S-LDSC)^40^ to assess how much of the heritability of gestational duration is contained within ATAC-seq, H3K4me1, H3K4me3, H3K27ac and pcHi-C peaks. LD scores were calculated using the peaks identified as reproducible across either treated or untreated samples as annotations and the 1000 Genomes Phase 3 European individuals (downloaded from the Price Lab website: https://data.broadinstitute.org/alkesgroup/LDSCORE/) as a reference LD panel, using only the HapMap3 SNP list (also from the Price Lab website). Stratified LD regression was performed on the gestational duration GWAS using the endometrial-tissue derived LD scores and the “baseline” LD scores contained in version 2.2 of the LD score regression baseline LD model. We include all annotations from the baseline LD model except those “flanking” annotations. This resulted in a total of 64 baseline annotations used in our S-LDSC analysis.

### Fine-mapping GWAS loci associated with gestational length

Fine mapping proceeded in three stages. In the first stage we partitioned the genome into 1,703 regions approximately independent regions using breakpoints derived by Berisa et al^39^. Next, we constructed a SNP-level prior probability that a particular SNP is causal. For this, we employed a Bayesian hierarchical model (TORUS^42^) that uses SNP-level annotations and GWAS summary statistics to estimate the extent to which SNPs with particular functional genomic annotations are more or less likely to be causal for a trait of interest. We ran TORUS with the gestational age GWAS summary statistics and the reproducible H3K27ac and H3K4me1 peaks from the treated samples along with the pcHi-C contact regions to obtain a SNP-level prior. Lastly, fine mapping was performed using a summary statistics-based version of the “Sum of Single Effects” model (SuSiE^43^), using 1000 Genome as reference panel. SuSiE (as implemented in the R package “susieR”) was run on the 10 regions believed to have one or more causal variants with FDR 0.1 as estimated by TORUS. For each region, SuSiE was run with a uniform prior (default setting of SuSiE) and with an informed prior learned by TORUS. The parameter L of SuSiE (maximum number of causal variants) is set at 1 when running SuSiE. This conservative setting ensures that the results are robust to possible LD mismatch between the reference panel and the GWAS samples. In fact, when L = 1, the PIP of a SNP depends only on its summary statistic (effect size and standard error), prior effect size and the number of SNPs in the locus, but not LD structure^65,66^.

### SNPs in Epigenome Roadmap histone modification peaks

H3K27ac, H3K4me1 and H4K4me3 histone modification peak coordinates were downloaded from https://egg2.wustl.edu/roadmap/data/byFileType/peaks/consolidated/narrowPeak/ucsc_compatible/ and bedtools intersect was used to find peaks that overlapped SNPs coordinates.

## Data availability

All the peaks sets, pcHi-C and gene expression data are available at https://www.immport.org/shared/study/SDY1626. Read count and TPM data are also available. An expanded set of 2,530 differentially expressed genes and sets of differential peaks called without statistically testing for fold-change is also made available. Source code for the GWAS enrichment analyses can be found at https://github.com/CreRecombinase/ptb_workflowr.

## Acknowledgements

The UChicago-Northwestern-Duke Prematurity Research Center was supported by a research grant from the March of Dimes to C.O., M.A.N., G.E.C., and J.K. This work was also supported by the March of Dimes Prematurity Research Center Ohio Collaborative and Bill and Melinda Gates Foundation (OPP1113966) to L.J.M. and G.Z. The authors acknowledge Christine Billstrand, Katie Naughton, Marcus Soliai and Raluca Nicolae for assistance with sample processing; Rabab Nasim, Sireesha Chinthala, and Rachel Loth for help with sample collection; Ruby Minhas for help with clinical questions and data entry; and Brian Furner and Seong Choi for designing and maintaining the sample tracking database. ALSPAC GWAS data: We are extremely grateful to all the families who took part in this study, the midwives for their help in recruiting them, and the whole ALSPAC team, which includes interviewers, computer and laboratory technicians, clerical workers, research scientists, volunteers, managers, receptionists and nurses. The UK Medical Research Council and Wellcome (Grant ref: 217065/Z/19/Z) and the University of Bristol provide core support for ALSPAC. A comprehensive list of grants funding is available on the ALSPAC website (http://www.bristol.ac.uk/alspac/external/documents/grant-acknowledgements.pdf). This research was specifically funded by Wellcome Trust WT088806 (Maternal genotype). This publication is the work of the authors and Xin He, Carole Ober and Marcelo A. Nobrega will serve as guarantors for the contents of this paper.

## Author contributions

C.O., M.A.N., G.E.C., and X.H designed and conceptualized the work. N.S., N.K. and J.M. analyzed data and interpreted results. I.A. coordinated all experiments. N.C., D.R.S., C.P., C.H., R.Z. and., H.K., and I.A. performed experiments, R.A. and S.R. facilitated sample collection. G.Z., B.J., M.H., K.T. and L.J.M. contributed the GWAS data. X.L., V.C.C., J.K., S.R., W.G., A.M., C.G. and I.A. contributed to discussions on study design. S.R., G.Z., L.J.M., V.J.L., G.E.C., C.O., X.H., and M.A.N. supervised aspects of the study. N.S., I.A., C.O., G.C., X.H. and M.A.N. wrote the manuscript. All authors read and commented on the manuscript.

